# Exploring the principles of embryonic mammary gland branching morphogenesis

**DOI:** 10.1101/2022.08.23.504958

**Authors:** Riitta Lindström, Jyoti P. Satta, Satu-Marja Myllymäki, Qiang Lan, Ewelina Trela, Renata Prunskaite-Hyyryläinen, Beata Kaczyńska, Maria Voutilainen, Satu Kuure, Seppo J. Vainio, Marja L. Mikkola

## Abstract

Branching morphogenesis is a characteristic feature of many essential organs such as the lung, kidney, and most glands, and the net result of two tissue behaviors: branch point initiation and elongation. Each branched organ has a distinct architecture customized to its physiological function, but how patterning occurs in these ramified tubular structures is a fundamental problem of development. Here we use quantitative 3D morphometrics, time-lapse imaging, manipulation of *ex vivo* cultured embryonic organs, and mice deficient in the planar cell polarity component *Vangl2* to address this question in the developing mammary gland. Our results show that the embryonic epithelial trees are highly complex in topology owing to the flexible use of two distinct modes of branch point initiation: lateral branching and tip bifurcation. This non-stereotypy was contrasted by the remarkably constant average branch frequency indicating a ductal growth-invariant, yet stochastic propensity to branch. The probability to branch was malleable and could be tuned by manipulating the Fgf10 and Tgf-β1 pathways. Finally, our *in vivo* and *ex vivo* time-lapse imaging suggested the involvement of tissue rearrangements in mammary branch elongation.

## Introduction

Branching morphogenesis is a fundamental developmental process driving the formation of a number of organs and tissues such as the mammary gland, salivary gland, kidney, lung, blood vessels, and nervous system (1, 2, 3). Branching involves repeated events of branch point formation and tube elongation enabling the maximization of the functional area of an organ in a limited three- dimensional (3D) space. Two distinct mechanisms of branch point formation have been described: tip splitting (or clefting) and side (lateral) branching (1, 4). Each organ has its unique way of generating the branching pattern – choice of branch site initiation, and length, diameter and spacing of branches – that gives rise to an organ-typical structure adapted for its specific function. For instance, lung branching morphogenesis is stereotyped. It involves three geometrically distinct modes of branching: domain (lateral) branching, planar bifurcation and orthogonal bifurcation.

Each of these modes occurs in a fixed order, thus lung branching is strictly regulated via pre- determined sequential branching events (5). Similar to lungs, kidney branching morphogenesis is highly reproducible (6). The ureteric bud undergoes a series of reiterative rounds of terminal bifurcations early and trifurcations later resulting in a ureteric tree where the extent and gross pattern of branch elaboration is conserved (7). Salivary gland morphogenesis also proceeds without side branching events and instead involves repetitive clefting of existing buds that rather widen than elongate before the next round of clefting events (8).

In contrast to many other organs, the branching of the mammary gland appears stochastic (9,10). It is also unique in that it generates an “open” architecture, a relatively sparse ductal network that leaves space for pregnancy-induced milk-producing alveoli. How the macroscopic features of the mammary gland including the size, network topology and spatial patterning, are designated remains poorly understood. Branching of the mammary gland takes place in distinct developmental stages each with discernible features: embryonic/pre-pubertal, pubertal, and reproductive (9). In mice, five pairs of small ductal trees with 10-20 branches form prior to birth by unknown mechanisms (11).

Initially, mammary rudiments branch as a solid mass of cells that soon after birth resolve into a bilayer of inner luminal and outer basal cells. Thereafter, the ductal system grows isometrically until the onset of puberty when ductal tips turn into larger, multi-layered bulbous terminal end buds (TEBs) that orchestrate the ensuing burst of ductal growth (12). 2D morphometric studies of fixed samples together with mathematical modeling suggest that branching during puberty is mainly driven by TEB bifurcations, while lateral branches emerge from previously formed ducts during the estrous cycle and pregnancy (10, 13).

Branch pattern formation is governed by reciprocal tissue crosstalk between the epithelial tree and the surrounding mesenchyme mediated by both shared and organ-specific secreted signaling molecules and extracellular matrix (ECM) components (2, 3). Embryonic mammary gland branching occurs in a mesenchymal fat pad precursor tissue that matures postnatally into a complex stroma consisting of adipocytes, fibroblasts, immune, and endothelial cells – all with established roles in morphogenesis (12, 14). Mammary gland development during puberty and reproductive life critically depends on systemic hormonal cues whose effects are, however, mediated by locally produced paracrine and autocrine factors thereby paralleling embryonic branching morphogenesis (9, 13). Many of these factors have been recognized including fibroblast growth factors (Fgf), Wnt ligands, transforming growth factor β1 (Tgf-β1), and epidermal growth factor (Egf) and tumor necrosis factor (Tnf) family members (9, 12, 15, 16, 17). However, what is less well understood is to what extent these pathways regulate growth and/or differentiation versus branch patterning *per se*. For example, while epithelial Fgf receptor signaling is pivotal for ductal branching, it is also essential for TEB maintenance and mammary epithelial cell self-renewal (18, 19, 20). The ligands involved are less well-defined. Fgf10 is the main ligand expressed in the mammary stroma (21), but assessing its role in *vivo* has been challenging as the germline deletion results in the absence of most mammary buds, and the mice do not survive beyond birth (22). Pathways inhibiting mammary gland development have also been identified, Tgf-β1 being the major one (23). Studies on postnatal mammary glands indicate that Tgf-β1 controls branching mainly by suppressing proliferation (24). On the other hand, *in vitro* studies suggest that branches might initiate at sites with a local minimum concentration of Tgf-β1 implying a specific role in branch patterning (25).

Much of our current understanding of lung, kidney, and salivary gland branching morphogenesis stems from studies utilizing established *ex vivo* organ cultures (2, 3, 6, 8). Tools to study mammary ductal growth and branching have been limited because the commonly used system in mammary gland research, the 3D organoid culture, is typically devoid of stromal cells and hence the critical epithelial-stromal crosstalk (26). Some recent organoid cultures do include stromal cells, yet their composition may not fully recapitulate that observed *in vivo* (27). Though numerous mouse studies have reported pubertal phenotypes, embryonic branching morphogenesis has remained a somewhat neglected area of research, possibly due to challenges in working with small mammary rudiments and/or the early developmental arrest observed in mice deficient in some key pathways such as Fgf10/Fgfr2b, and Wnt (11, 22, 28).

In this study, we aimed to decipher the principles of branching pattern formation in the mammary gland by combining 3D morphometrics with *ex vivo* embryonic mammary gland culture method that preserves the epithelial-mesenchymal tissue interactions intact (29). Our results show that the topologies of embryonic mammary glands are highly variable suggesting a stochastic branching process, yet the average branching frequency remains remarkably constant throughout development and thereby ‘stereotypic’. Branch point formation, however, was sensitive to exogenous manipulation of Fgf10 and Tgf-β pathways. Time-lapse imaging revealed that lateral branching and tip bifurcation jointly drive embryonic mammary gland morphogenesis, a finding supported by the 3D optical projection tomography data. Our data also imply that tissue-level rearrangements may contribute to ductal elongation.

## Results

### Embryonic mammary gland branches stochastically, yet at a constant rate

To characterize the branching of embryonic mammary glands in 3D, fixed mammary glands were collected and scanned with optical projection tomography (OPT). Samples were dissected at embryonic day (E) 16.5 (when the branching morphogenesis commenced in most anterior glands), E17.5, and prior to birth at E18.5 from mice of mixed background. In total, 101 samples of mammary glands 1 to 5 ranging from unbranched mammary primordia (MG5 at E16.5) to glands with up to 25 ductal tips (MG1 at E18.5) were analyzed (Fig 1A and S1A Fig). As previously suggested (30), the anterior mammary primordia were noticeably larger than the posterior ones (Fig 1A and S1 Movie), and the patterns of ramifications differed from embryo to embryo, as well as between the left and right side of the same embryo (S1B Fig). Already at E16.5, the 3D structure of the mammary gland was relatively flat reflecting its growth between the skin and muscle tissue (Fig 1B, S1 Movie).

**Fig 1.**
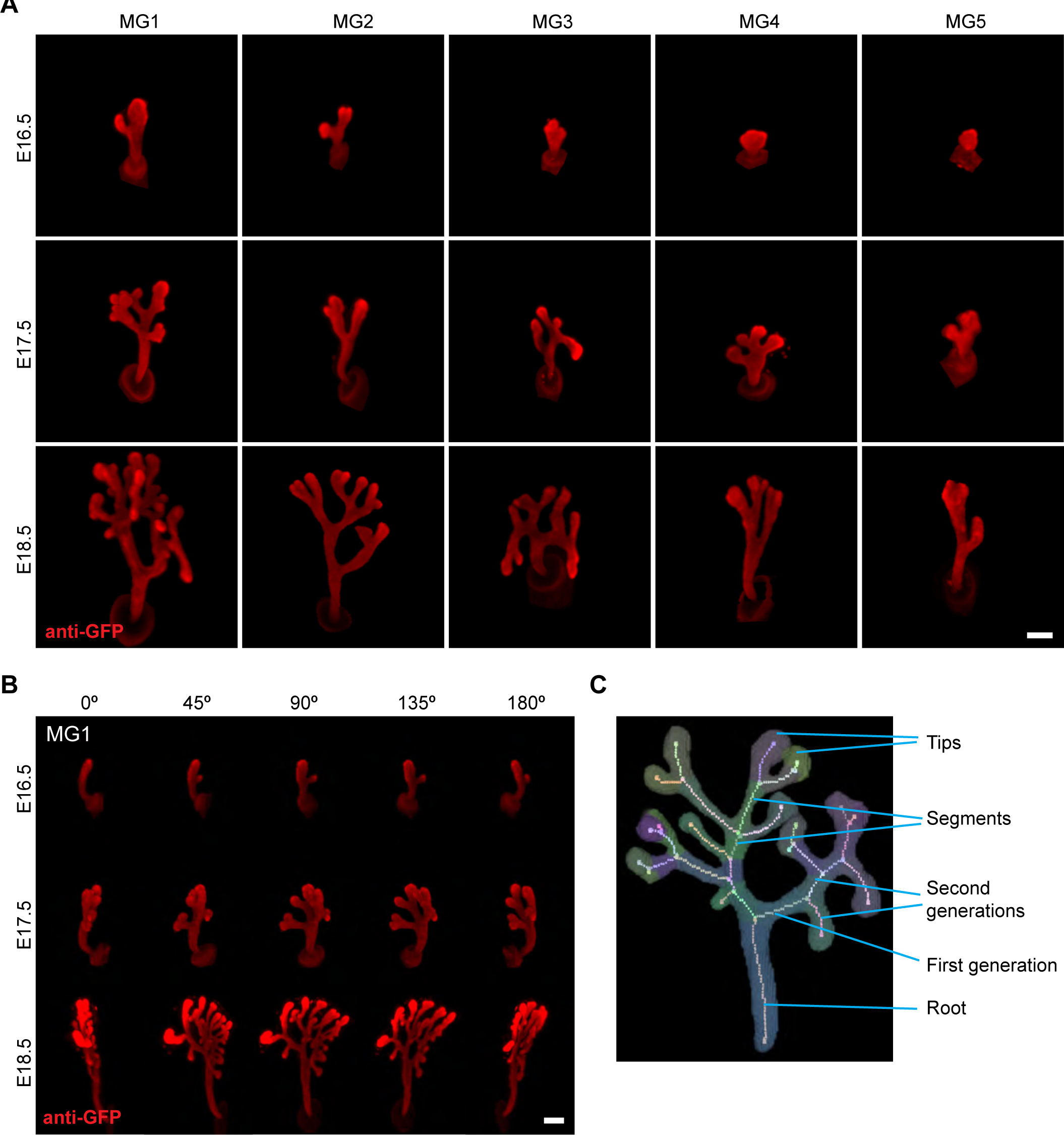
Embryonic mammary glands develop asynchronously. **(A)** Representative OPT images of K14-Cre;mT/mG mammary glands (MG) 1 to 5 stained with anti-GFP antibody at E16.5, E17.5, and E18.5. **(B)** 180° rotational views of mammary gland 1 at E16.5, E17.5, and E18.5. **(C)** The terminology used throughout the manuscript; the colors highlight different segments. Three types of branches including the three branch types (root, segment, and tip) are indicated. The root is the main duct between the nipple and the first branch point, the segment is an ‘internal branch’ between two branch points, and the tips are terminal branches. Each segment is assigned a generation number relative to the root. Scale bar is 150 µm.

For quantitative spatial analysis, images were analyzed with the Tree Surveyor software that enables derivation of network properties such as tip number, branch length and angle, tree volume, and branching generations (31) (S1C Fig). Quantification confirmed that branching increased from E16.5 to E18.5 based on both tip and generation numbers (Fig 2A and 2B). In principle, the generation number can be used to infer the sequence of branching events, counting from the root of the ductal tree (for terminology used in the study, see Fig 1C). When tip number was plotted against the number of generations per gland, it was evident that tips were not propagated as expected in a stereotypic bifurcating system where each terminal branch gives rise to two new ones (S1D Fig).

**Fig 2.**
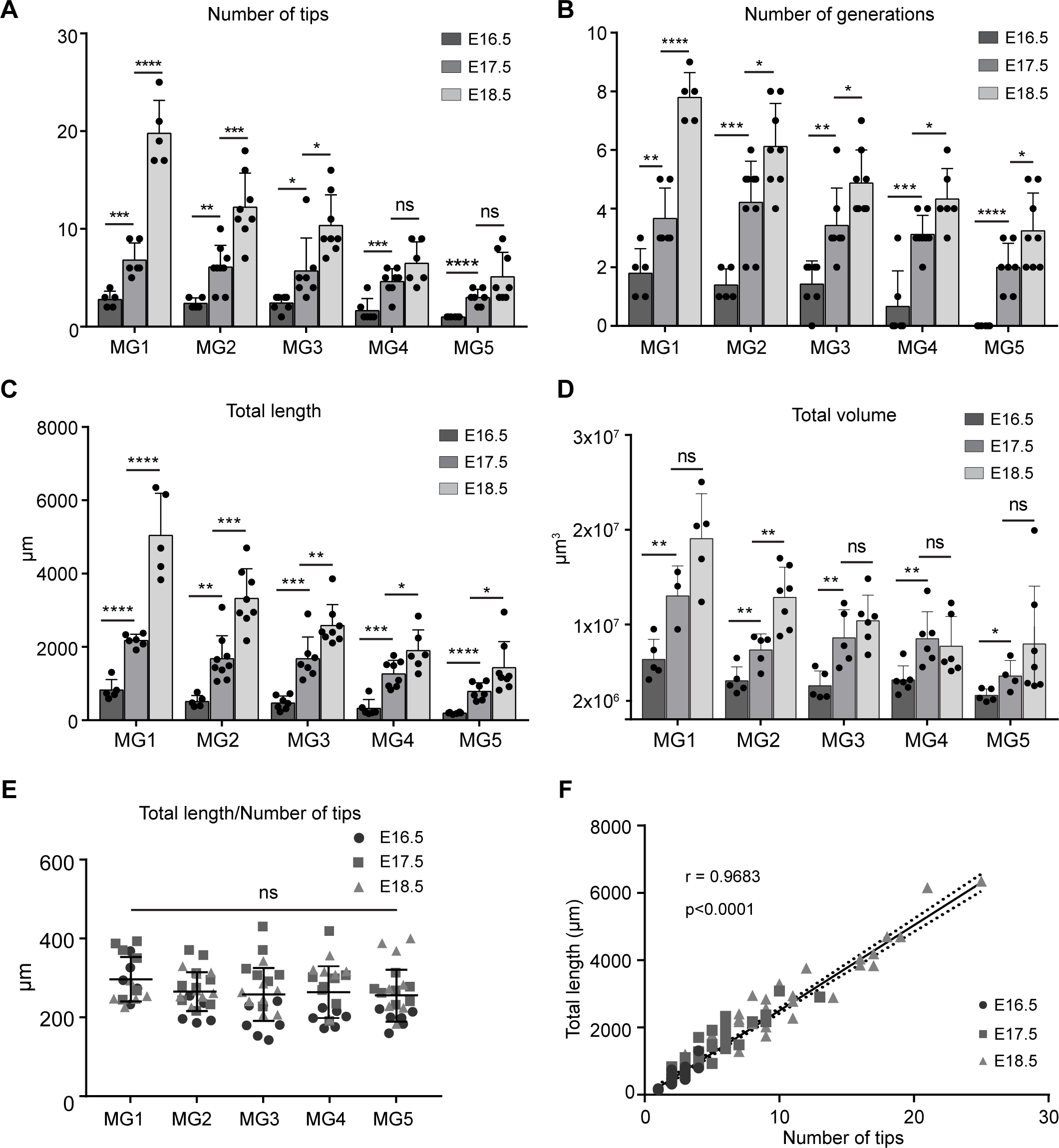
Embryonic mammary glands branch at a constant frequency. **(A-D)** Quantification of the number of ductal tips **(A)** and branch generations **(B)**, as well as the total length **(C)** and total volume **(D)** of the ductal tree in mammary glands (MG) 1-5. Student’s t-test was used to assess statistical significance. **(E)** The ratio of total length to tip number is constant over developmental stages and across mammary glands 1-5. Statistical significance was tested with one-way ANOVA. Data in (A-E) are shown as mean ± SD. **(F)** The correlation of the total ductal length and tip number of all mammary glands was assessed with Pearson’s r.

Quantification of the total length and volume also confirmed the increase in size in each gland between E16.5 and E18.5, again the anterior glands being larger than the posterior glands (Fig 2C and 2D). However, the ratio of total ductal length and the number of ductal tips showed no overt difference across glands (Fig 2E) indicating that despite the difference in size, the branch point frequency is constant, and hence each gland branches essentially the same way. Strikingly, when all glands were pooled for analysis, the total length of the ductal tree and the number of ductal tips showed a near-perfect positive correlation (Pearson’s r=0.968) (Fig 2F). On average, there was one tip for every ∼260 µm ± 40 µm (mean ±SD) of ductal length, a value that was constant over the developmental stage of the mammary glands (using tip number as a proxy for developmental stage), although less developed glands showed much higher variation compared to more advanced ones (S1E Fig). A strong positive correlation was also observed between the total volume and number of tips (Pearson’s r=0.8629) (S1F Fig).

Taken together, our OPT images show that each embryonic mammary gland displays a unique non- stereotypic branching pattern. Remarkaby, however, they displayed a steady average increase in branch point frequency.

### Both lateral branching and terminal bifurcations drive pre-pubertal mammary gland branching

As shown above (Fig. S1D Fig), the topology of embryonic mammary glands did not confine to a stereotypic bifurcating system. This can be either because some terminal branches cease to grow, as previously suggested to take place during puberty (10) or alternatively, branches may form by side branching which changes the generation dynamics, as a generation would be divided in two “retrospectively”. Since it is not possible to distinguish between these two options based on fixed samples, we utilized an in-house developed organ culture system to perform time-lapse imaging of *ex vivo* cultured embryonic mammary glands. This culture set-up recapitulates growth *in vivo* relatively faithfully (29, 32). E13.5 mammary buds expressing epithelial GFP (K14-Cre; mT/mG) were dissected with the surrounding mesenchyme, and imaging was started on day three of culture when branching had been initiated. Images were captured every four hours for up to four days (Fig 3A, S2 Movie). Time-lapse imaging revealed that lateral branch formation was the main mode of branching, covering ∼75% of all branching events (Fig 3B and 3C). Of these, 63% formed in the tips (i.e. in terminal branches; Fig. 3B upper panel shows an example of a lateral branch forming in a tip) and 37% in segments (i.e. between two existing branch points). All glands monitored had bifurcation events (24% of branching events), as well. Terminal trifurcations were also observed but rarely (<2% of all events).

**Fig 3.**
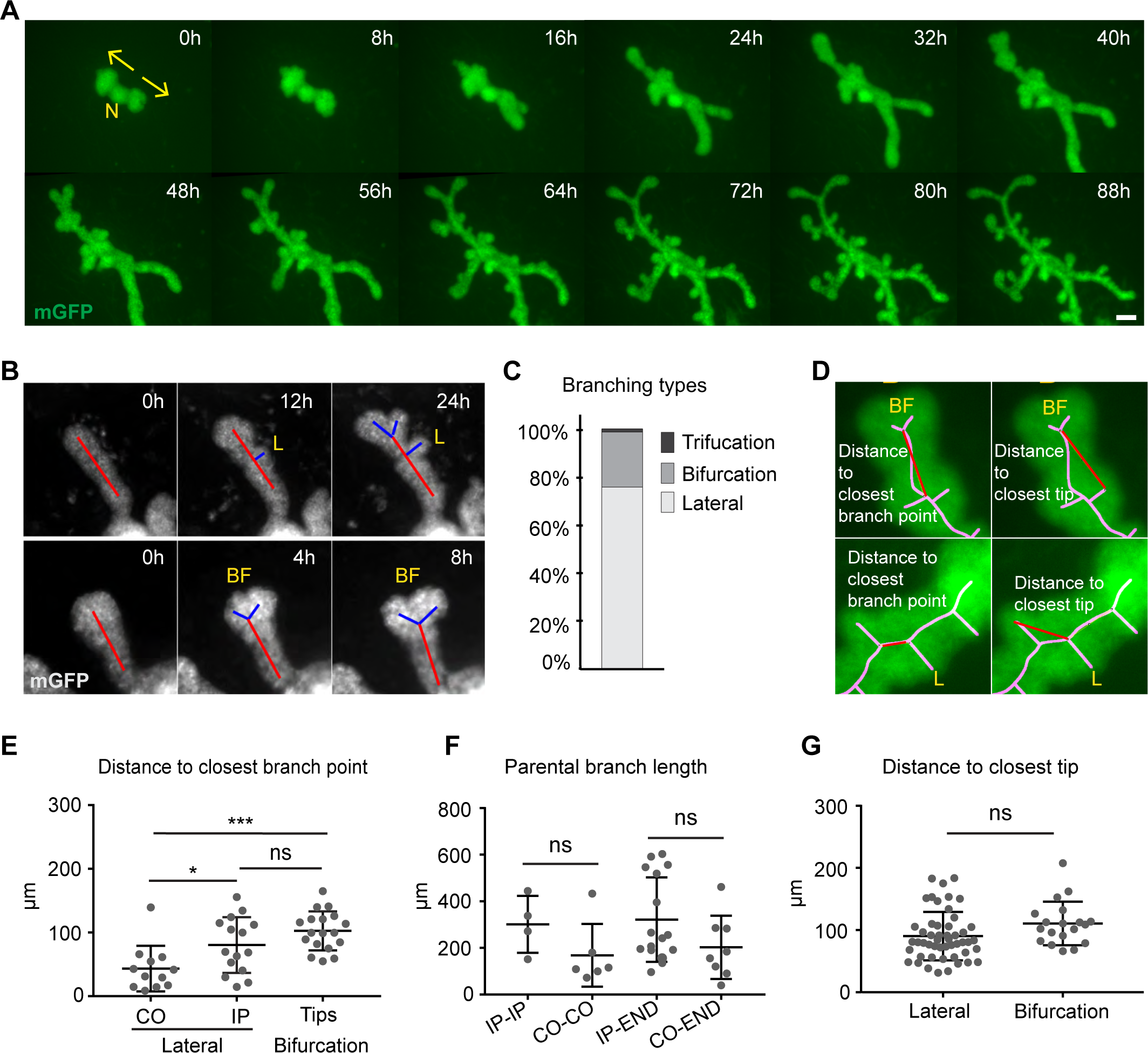
Embryonic mammary gland branches via side branching and terminal bifurcations. **(A)** Time-lapse series of *ex vivo* cultured K14-Cre;mT/mG mammary glands. Mammary buds were dissected at E13.5, imaging was started after 3 days of culture and continued for up to 96 hours. N, nipple; arrows show the direction of growth. **(B)** An example of a time-lapse video used to assess and quantify the types of branching events: lateral branching (L), and bi- and trifurcations (BF). **(C)** Quantification of the types of branching events (n=149 branching events from 15 mammary glands). **(D)** Images depict how the branch point and tip closest to an emerging bifurcation (top) and lateral branch (below) were quantified. **(E)** Quantification of the distance to the closest branch point of nascent lateral branches and branches formed by bifurcation. For lateral branches, contra- (CO) and ipsilateral (IP) branch points were analyzed separately. **(F)** Quantification of the parental branch length at the time when a new lateral branch emerged between two ipsilateral branches (IP-IP), between two contralateral branches (CO-CO), between a terminal branch and an ipsilateral side branch (END-IP), and between a terminal branch and a contralateral side branch (END-CO). **(G)** Quantification of the distance to the closest branch tip of nascent lateral branches or branches formed by bifurcation. Data in (D-F) are shown as mean ± SD. Student’s t-test was used to assess statistical significance. Scale bar is 100 µm.

As most explants had already started to branch when we initiated our imaging, we could not analyze the type of the very first branching event. To assess this indirectly, we turned to our OPT dataset and compared the length of the root (i.e. main duct) in mammary sprouts that had not yet branched to those that had branched just once. Quantification revealed that the length of the root was 40% shorter and the volume 54% smaller in rudiments with two tips compared to unbranched ones (S2A and S2B Fig), a finding poorly compatible with a tip splitting event. A pairwise comparison of the tip lengths in glands with two tips only showed that while in some glands they were of equal length, others displayed a notable difference (S2C Fig). In conclusion, our live imaging data establishes that pre-pubertal branching morphogenesis utilizes a combination of lateral branching and terminal bifurcation events. Fixed sample data imply that the mode of the very first branching event is also stochastic.

### Neighboring tips constrain the formation of new branches

Having identified both lateral and terminal branching events, we next analyzed in detail where they occur within the epithelial network and if their locations follow any trend. For this purpose, we measured the distance of incipient lateral and bifurcating terminal branches to their closest neighboring tips and branch points (Fig 3D). Tip splitting occurred at a minimum distance of 55 µm from the last branch point (Fig 3E), indicating that terminal branches need to elongate to a certain length and/or grow to a certain size before bifurcating. In contrast, the occurrence of lateral branches was not as limited by their distance to existing branch points, although it did make a difference at which side other branches were. Branch points on the opposite sides (contralateral, CO) were tolerated at closer distances (minimum 9 µm; average 44 µm) than branch points on the same side (ipsilateral, IP) (minimum 15 µm; average 80 µm), possibly due to the fact that branching is constrained in the Z-axis both *in vivo* and *ex vivo* (Fig 3E). Accordingly, the length of the parental branch was dependent on the orientation of the closest branches. When a new branch emerged between two existing branches, the segment was longer between two ipsilateral branches (IP-IP) compared to two contralateral branches (CO-CO) although the difference did not reach statistical significance (Fig 3F). The same was true when a lateral branch emerged in a terminal branch (END- CO vs. END-IP) (Fig 3F). Collectively, these data suggest that rather than branch points as such, proximity to other tips might limit the location of a new lateral branch. Indeed, neighboring tips were at a minimum distance of 31 µm (mean 90 µm) from an emerging lateral branch; similar findings were made with bifurcations (Fig 3G). Thus, neighboring tips, rather than branch points (except for ipsilateral branches), spatially constrain the emergence of new branches.

### Spatial analysis of branch parameters predicts the pattern of branching, elongation, and growth

Previous studies on the pubertal mammary gland localized branching and elongation to terminal branches, indicating that the length of a branch (segment) is determined once the tip bifurcates and is preserved thereafter (10). To gain more insight into branch dynamics, we turned to our OPT dataset and analyzed separately terminal branches (tips), segments (branches between two branching points), and roots. Segments were found to be significantly shorter than terminal branches (Fig 4A), a finding that cannot be explained if branching occurs via tip bifurcation only. However, as shown by our time-lapse data (Fig 3B), both segments and tips could be shortened through lateral branching. The roots on the other hand were exceptionally long compared to both segments and terminal branches (Fig 4A), as well as thick and large as shown by the median diameter and volume, respectively (Fig 4B and 4C, see also below). To further investigate how growth contributes to branch elongation in different parts of the gland, we visualized the dimensions of terminal branches and segments as a distribution of their length to the median diameter (Fig 4D). The segments were narrower the longer they were, but the opposite was observed in terminal branches suggesting that they may widen before they branch. We also assessed whether there was any correlation between the median diameter of the branch and the developmental stage of the gland (using tip number as a proxy). The segments got narrower as the gland advanced in development, but the terminal branches behaved the opposite (Fig 4E), possibly owing to the fact that on average they were shorter in more advanced glands (S3A Fig).

**Fig 4.**
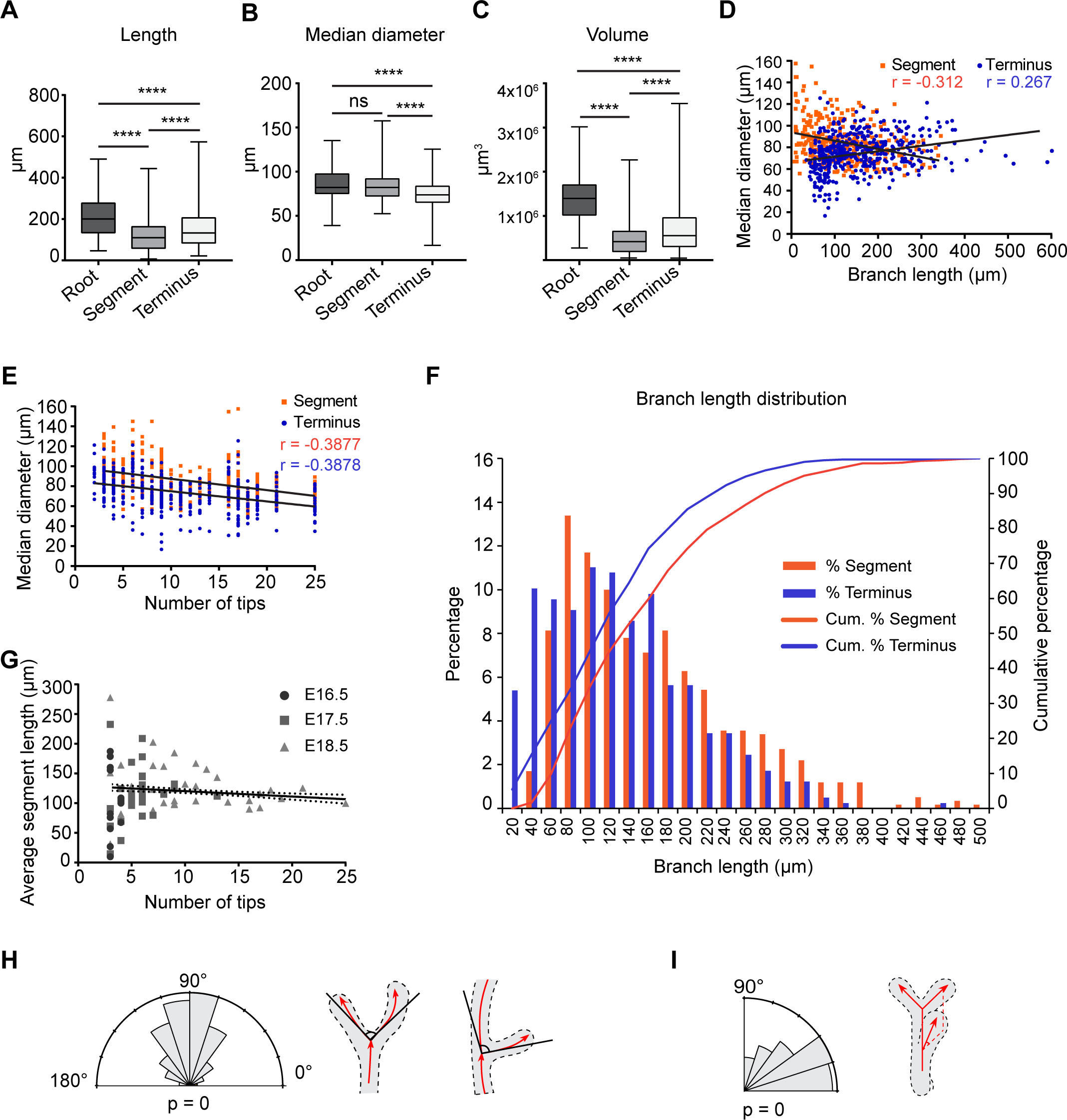
Branch dynamics in embryonic mammary glands. Branch parameters were quantified with Tree Surveyor. Quantification of the length of roots (n=89), segments (n =409) and terminal branches (n=592) **(A)**, median diameter of the length of roots (n =73), segments (n =344) and terminal branches (n=465) **(B)** and volume **(C)** of roots (n=73), segments (n=347), and terminal branches (n=496) from all mammary glands at all developmental stages. The data shown represent the median (line) with 25th and 75th percentiles (hinges) plus the min to max ranges (whiskers). The correlation of the median diameter to branch length **(D)** and the number of tips **(E)** of segments (n=344) and terminal branches (n=465) was assessed with Pearson’s r. **(F)** Length distributions of segments and terminal branches. **(G)** Correlation of the average segment length of each mammary gland to the number of tips (n=408) was assessed with Pearson’s r. **(H, I)** Schematics depicting the local “bifurcation” (n=491) (**H)** and dihedral (n=395) **(I)** branch angles and their quantifications. Rayleigh test for nonuniformity, H0=random. (#for root parameters, only branched glands were included).

Branch length measures elongation between two branching events that may or may not be sequential, as we have learned. In the pubertal mammary gland, the timing between consecutive branching events appears to be random (33).

To understand how the spacing of branching events is established, we analyzed the distribution of segments and terminal branch lengths, reflecting the probability to branch. The distribution of segments was skewed towards shorter lengths, indicating that there is a tendency to minimize spacing between two branch points (Fig 4F and S3B Fig). A notable fraction (∼25%) of all segments were shorter than 55 µm, the minimum distance from a bifurcating tip to the closest branch point (Fig 3D), a finding further supporting our conclusions on the contribution of side branching in mammary gland morphogenesis. Given that spatial constraints become greater the shorter the segment is, this minimum likely approaches the maximum epithelial packing density that is tolerated. Interestingly, when the distribution of the average segment length of each gland was analyzed against developmental time, we noticed much higher variation in the younger glands compared to the older ones (Fig 4G).

The branching angles also contribute to the topology of the epithelial network. Bifurcating TEBs branch with an average angle of 70-75 degrees thereby biasing the global orientation of the ductal tree along the long axis of the fat pad (34, 35). We found the local ‘bifurcation’ angles of branch points (the angle between the start direction of two branches) to be larger, focusing around 90 degrees (Fig 4H). This finding is in line with the notion that the longitudinal bias is not observed in 2-week-old, pre-pubertal mice (35), In contrast, rotation between consecutive branch points (dihedral angle) was small (Fig 4I), consistent with the rather 2-dimensional architecture of the mammary gland.

### Tgf-β1 and Fgf10 change branch point frequency

Mammary branching morphogenesis is regulated by conserved secreted signaling molecules, but their contribution to growth versus branch patterning is challenging to discern (4, 9, 12). Some may simply modulate growth, while others might affect the propensity to branch. Our *ex vivo* culture system provides a means to distinguish between these two options prompting us to assess how Tgf- β1 and Fgf10 regulate ductal morphogenesis. Since there is no evidence of long-range guidance cues in the mammary gland stroma, and *Fgf10* is uniformly expressed in the fat pad (18), we reasoned that application of the candidate factors in the growth medium would best mimic the *in vivo* situation. Mammary buds were isolated from E13.5 embryos expressing epithelial GFP, allowed to grow for 2 days before applying either the signaling molecules or their inhibitors, and then cultured for further 5 days. We quantified the total ductal length and the tip number and calculated their ratio to assess if branching frequency was affected.

To assess the impact of Tgf-β1 on branching rate, we opted for a low concentration given its well-known strong anti-proliferatory effect (23, 36). When Tgf-β1 (2.5 ng/ml) was administered to explant cultures, a significant reduction in the number of tips was observed, and fewer tips were generated per total ductal length indicating a decrease in branching frequency (Fig 5A and 5B).

**Fig. 5.**
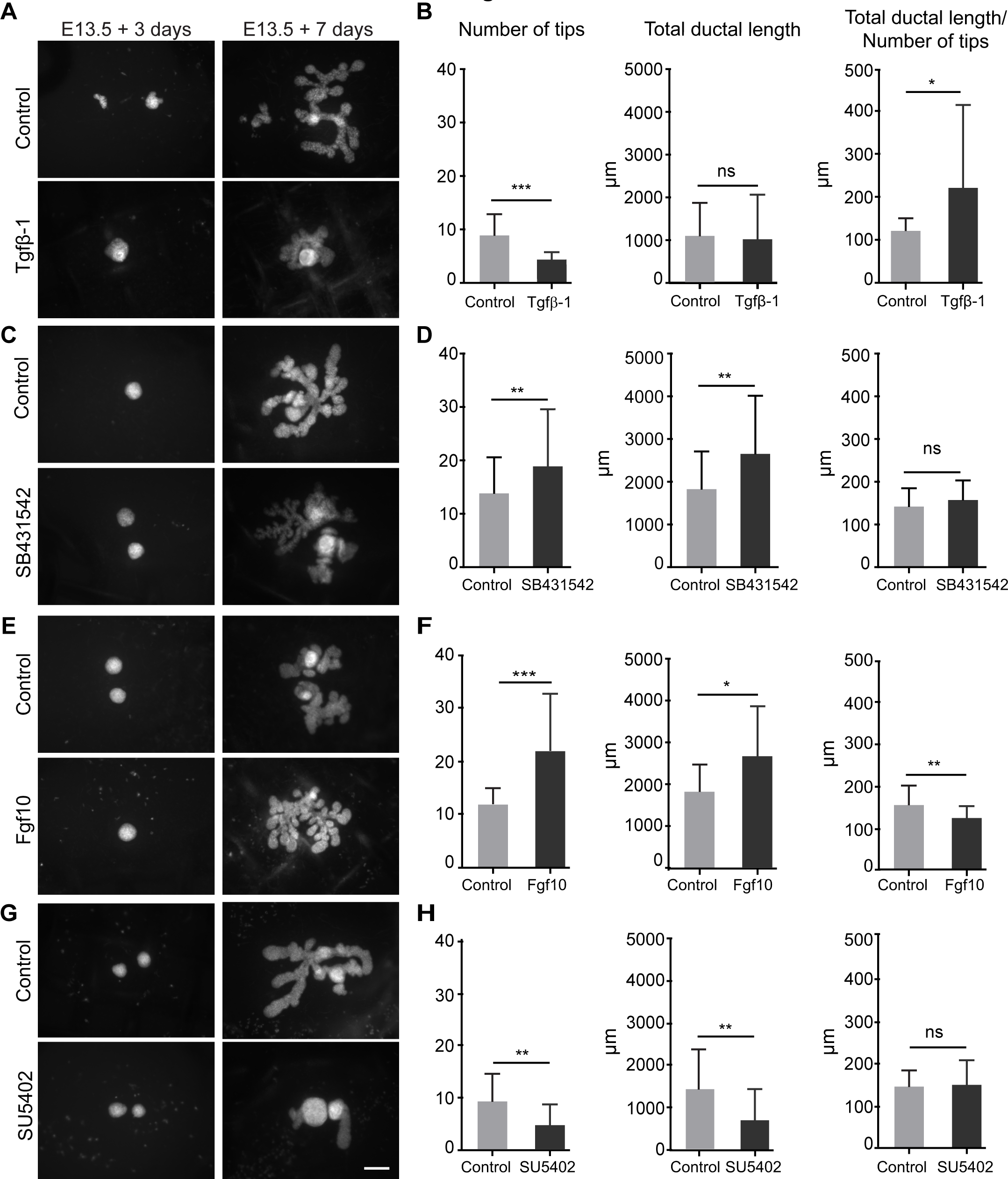
Ex vivo manipulation of cultured mammary glands with Tgf-β1 and Fgf10 pathways. **(A, C, E, G)** Mammary buds were dissected at E13.5, the indicated signaling molecules/pathway inhibitors were added on day 2 and the explants were cultured for further 5 days. **(B, D, F, H)** The number of tips and total ductal length and their ratio were quantified on day 7. **(A, B)** Tgf-β1 (2.5ng/ml) (n_ctrl_=15, n_Tgf-β1_=20 from 2 independent experiments). **(C, D)** SB431542 (10 µM) (n_ctrl_=60, n_SB431542_=65 from 8 independent experiments). **(E, F)** Fgf10 (100ng/ml) (n_ctrl_=21, n_Fgf10_=23 from 3 independent experiments). **(G, H)** SU5402 (2.5 µM) (n_ctrl_=24, n_SU5402_=21 from 3 independent experiments). Data are shown as mean ± SD. Statistical significance was assessed with Student’s t-test. Scale bar is 200 µm.

Treating explants with the Tgf-βRI inhibitor SB431542 had the opposite effect. The number of tips and total ductal length both increased, yet without a change in their ratio (Fig 5C and 5D). Curiously, the ducts also got narrower, for reasons that currently remain undefined. The addition of Fgf10 (100 ng/ml) to mammary gland explant cultures significantly increased both the number of ductal tips and total ductal length (Fig 5E and 5F) the latter indicating that is enhances growth, as expected. Also branching frequency was augmented as shown by the analysis of total ductal length per number of tips (Fig 5F). Conversely, the broad-spectrum Fgf receptor inhibitor SU5402 severely hindered growth as evidenced by the reduced total length of the ductal network and tip number though branch frequency remained unaffected (Fig 5G and 5H).

### Tissue rearrangements contribute to main duct elongation

Next, we wanted to better understand the process of branch elongation and first focused on the main duct (root), which we can easily identify in fixed glands by its connection to the skin. Unlike terminal branches and segments, root lengths conformed to a rather normal distribution (S3B and S3C Fig) suggesting that branching is prevented close to the skin. Embryonic stage-by-stage analysis revealed statistically significant elongation of the root (Fig 6A), a finding confirmed by plotting the root length against the developmental stage of the mammary gland (using the total tip number as a proxy) (Fig 6B). This result was surprising given that branch elongation has mainly been associated with the growing tips (10) and prompted us to examine cell proliferation in the root before and after the onset of branching morphogenesis. To this end, we analyzed fixed E16.5 mammary glands (ranging in size from non-branched ones to those with 5 tips) of bi-transgenic Fucci cell cycle indicator mice (37), where cells express nuclear green (mAzami Green, mAG) and red (mKusabira Orange, mKO2) in S/G2/M and G1/G0 phases of the cell cycle, respectively (Fig 6C). Most cells in the root were quiescent, regardless of the developmental stage, while the tips were proliferative, as expected (Fig 6D), suggesting that mechanisms other than proliferation drive root elongation.

**Fig 6.**
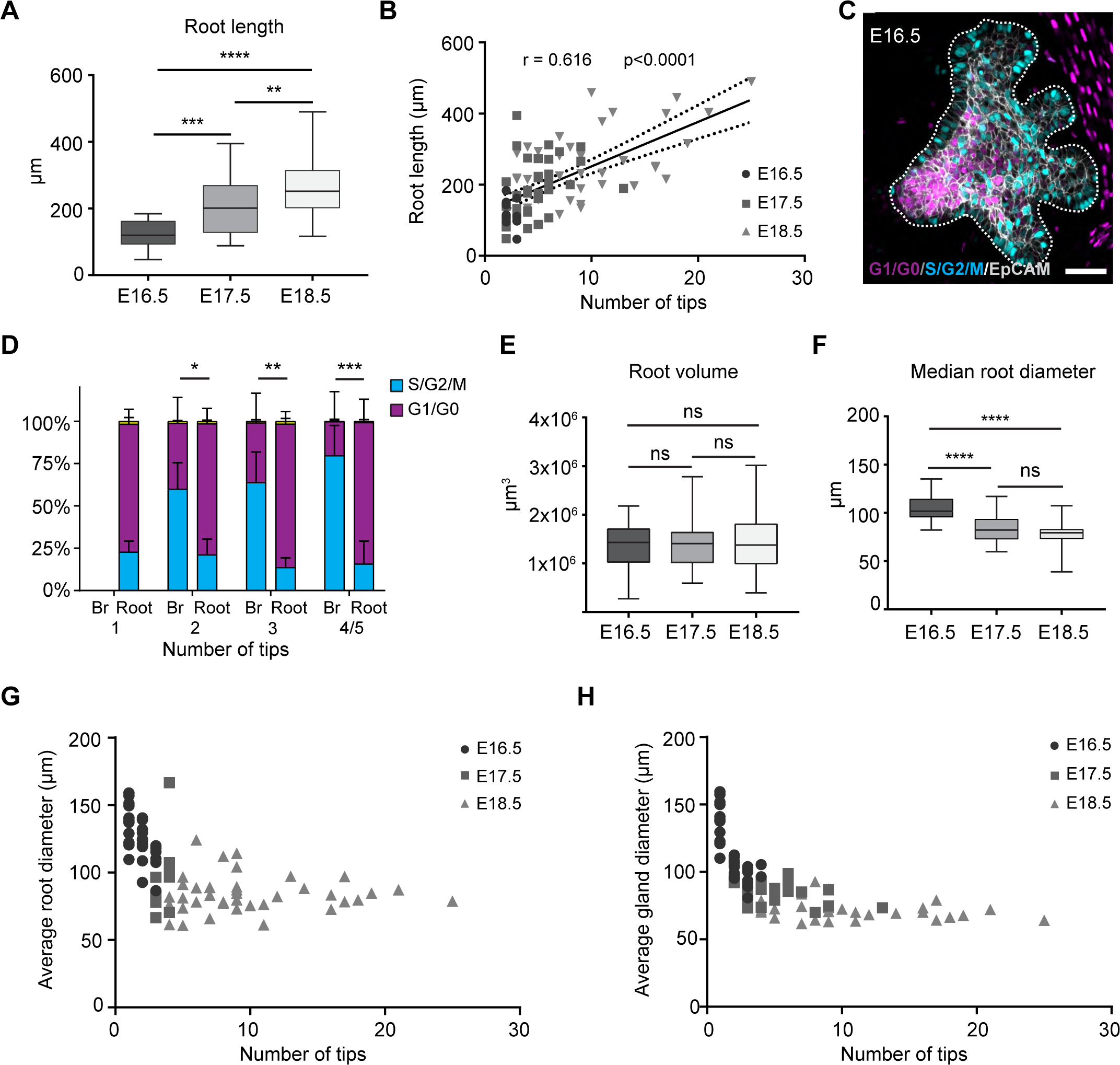
The main duct elongates and narrows as development advances. **(A)** Quantification of root length from mammary glands 1-5 at E16.5 (n=18), E17.5 (n=36), and E18.5 (n=35). **(B)** Correlation of the root length to the number of tips of all glands was assessed with Pearson’s r. **(C)** A representative image of an E16.5 MG3 from Fucci mice, cells in S/G2/M in cyan, and cells in G1/G0 in purple. The dashed line shows the epithelial-mesenchymal border identified by EpCAM immunostaining. **(D)** Quantification of the proportions of cells in G1/G0 and S/G2/M (or expressing both markers, in yellow) in roots and branches (Br) in mammary glands with one (unbranched) (n=2), two (n=3), three (n=4) or four to five tips (n=5). Statistical significance was assessed with Student’s t-test. **(E)** Quantification of root length from mammary glands 1-5 at E16.5, E17.5, and E18.5. **(F)** Quantification of median root diameter from mammary glands 1-5 at E16.5, E17.5, and E18.5. (#for root parameters, only branched glands were included) **(G)** Average root diameter plotted against the number of tips (n=84) reveals that the diameter gradually settles ∼70 µm. **(H)** Average gland diameter plotted against the number of tips (n=71) reveals that the diameter gradually settles ∼70 µm.

Next, we analyzed the root volume and diameter at E16.5-E18.5. These data clearly show that while the root elongates, it does not grow in volume, but narrows in diameter (Fig 6E-G). Together, these data show that the root elongates not by increasing its volume, but mainly by changing its aspect ratio from shorter and thicker to longer and thinner. The average diameter of, not only the root but also the entire gland narrowed during branching morphogenesis settling down at ∼70 µm (Fig 6G, H; see also Fig 4E), a value very close to that of mature ducts in pubertal glands (34).

Unfortunately, for reasons that are not clear to us, the elongation of the root was rarely observed in our *ex vivo* cultures precluding analysis by time-lapse imaging. Instead, we asked whether a similar process occurs elsewhere in the gland, and assessed if segments could also narrow down and elongate even ‘behind’ a branching event (S3D Fig). To this end, we measured the dimensions of newly formed segments from the time-lapse videos. Indeed, a significant change was observed in the aspect ratio: the segments elongated while their diameter got narrower (S3E and S3F Fig).

### Loss of Vangl2 compromises growth yet accelerates branching

Narrowing of the root without a change in the volume points to convergent extension, a mechanism where the tissue converges (narrows) along one axis to lengthen it on a perpendicular axis, either via cellular movements or coordinated cell shape changes (38). Convergent extension is mediated through the conserved planar cell polarity (PCP) pathway, which in mammals is established upon asymmetrically distributed Van Gogh-like (Vangl) membrane proteins, Vangl1 and Vangl2. To ask if main duct elongation is regulated by PCP/Vangl2, we collected *Vangl2* null and wild type control C57Bl/6 mammary glands at E17.5 when branching had started in all glands and the main duct was elongating, and used 3D confocal microscopy for analysis (Fig 7A and S4A). Again, anterior glands were larger than posterior ones, though in C57Bl/6 background MG2 was the most advanced one (S4A and S4B Fig). The phenotype of *Vangl2* null mammary glands appeared variable (Fig 7A and S4A), verified by the quantifications of tip number and total volume, and best exemplified by MG1- 3 (S4B and S4C Fig). For instance, the number of tips was not affected except in MG1: *Vangl2* mutants had significantly more tips, yet there was no change in the volume. On the other hand, MG2 and MG3 of *Vangl2* mutants were greatly reduced in size although the number of tips was not altered (S4B and S4C Fig). Together these findings suggest that Vangl2 may regulate the rate of branching. To test this hypothesis, we compared control and *Vangl2* mutant mammary glands of equal tip numbers. The volume of *Vangl2* mutant glands was consistently smaller (Fig 7B) and hence the tip number per volume of the gland was significantly (∼60%) higher in *Vangl2* mutants compared to their wild type littermates (Fig 7C) indicating higher propensity of branch point formation despite overall growth impairment. Analysis of roots revealed that on average they were 35% smaller in volume, 26% shorter in length, but only 7% smaller in average diameter (Fig 7D-F and S4D-F). Normalization of the root volume, length and average diameter to tip number confirmed that mainly the volume and length were affected, not the diameter (S4D-F Fig).

**Fig 7.**
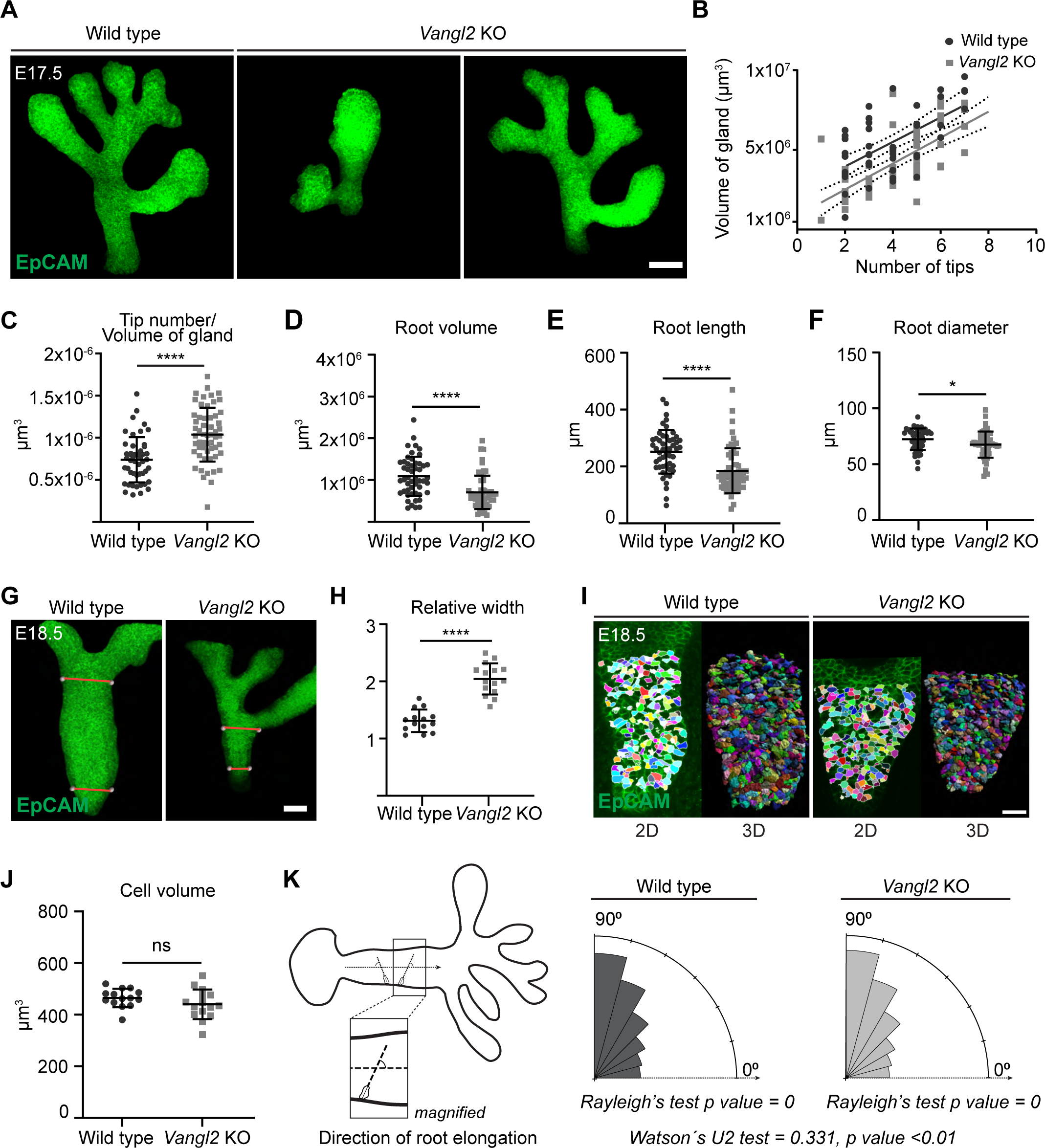
Loss of Vangl2 compromises mammary gland growth but accelerates branching. **(A)** Representative confocal maximum intensity projection images of EpCAM-stained E17.5 mammary gland (MG) 2 of wild type control (*Vangl2^+/+^*) and two *Vangl2* knockout (KO) embryos. Scale bar is 100 µm. **(B)** Correlation of the mammary gland to number of tips in wild type littermates and *Vangl2* KO embryos was assessed with Pearson’s r (n_wt_=49 and n*_Vangl2_*_KO_=57). **(C)** Quantification of the ratio of number of ductal tips to the volume of glands of mammary glands 1-5 in wild type littermates and *Vangl2* KO embryos. **(D-F)** Root volume **(D)**, root length **(E)**, and root diameter **(F)** of mammary glands 1-5 in wild type and *Vangl2* KO embryos. **(G)** Representative confocal maximum intensity projection images of EpCAM-stained E18.5 mammary gland (MG) of wild type control (*Vangl2*^+/+^) and *Vangl2* knockout (KO) embryos. Scale bar 50 µm. **(H)** Quantification of relative width of the root measured near the first branch point/close to the nipple in wild type and *Vangl2* KO embryos (n_wt_=14 and n*_Vangl2_*_KO_=15). **(I)** Cell segmentation based on EpCAM in the wild type and *Vangl2* KO mammary glands. Scale bar 30 µm. **(J)** Quantification of the cell volumes in wild type (n=14) and *Vangl2* KO (n=15) main ducts. **(K)** A schematic showing the cell orientation analysis in the main duct (root), and Rosa plots of cell orientation angles with respect to the elongating main duct; n_wt_=10766 cells, n*_Vangl2_*_KO_= 22008 cells. Statistical significance was assessed with Student’s t-test (B-J), and Watson’s U2 test, and Rayleigh’s test for nonuniformity, H0 = random (K).

To gain deeper insights into the role of Vangl2 in main duct elongation, we focused on embryonic stage E18.5, the latest stage we could obtain live mutant embryos. Mammary glands were stained with EpCAM and phalloidin, and the main ducts were imaged at high resolution in 3D. Again, the main ducts of *Vangl2* mutants were abnormally short and in contrast to wild types, often “V- shaped”: narrower near the nipple and wider near the first branch point (Fig 7G), a finding confirmed by quantification (Fig 7H). As two distinct processes take place during late embryogenesis, lumen formation and ductal elongation, and might affect one another, we first analyzed the former. In both control and *Vangl2* mutants formation of microlumini was evident and associated with high levels of F-actin suggesting no major impairment in lumen formation (S4G Fig). For cell level analysis, cells were segmented in 3D based on EpCAM staining (Fig 7I). There was no difference in cell volumes between the two genotypes (Fig 7J).

Convergent extension can be either passive (the surrounding environment deforms the tissue) or active (cells of the elongating tissue generate the necessary forces) (39). Cell intercalation takes place in both, but typically the axis of cell elongation is parallel to the axis of tissue elongation in the former, and perpendicular in the latter. Quantification of cells’ longest axis with respect to the axis of the elongating main duct revealed that cells in wild type main duct were non-randomly distributed, and preferentially oriented perpendicular to the long axis of the main duct (Fig 7K: 0° equals to perfect alignment with the elongation axis). The same was true in *Vangl2* mutants.

However, there was a statistically significant difference in the cell orientation between the two genotypes.

## Discussion

In this study, we took advantage of the Tree Surveyor software designed to interrogate branching morphogenesis in 3D (31) and *ex vivo* time-lapse imaging (29) to provide a comprehensive description of the embryonic mammary gland branching morphogenesis by mapping and quantifying the embryonic ductal network and its dynamic changes over time. The topologies of the individual epithelial trees were highly variable, as reported for pubertal and adult glands (10, 40), evidencing that mammary branching is stochastic from the very beginning. Our findings are in line with the idea of mammary branching morphogenesis as ‘random branching and exploratory walk’ ultimately leading to efficient filling of the available space (33, 35). In pubertal glands, the model entails termination of branches due to crowding, i.e. when tips arrive too close to another duct, tip, or the border of the fat pad (33). However, such annihilation behavior is likely to be less relevant (or non-existing) in the embryo where the stroma can readily accommodate much larger ductal trees, as exemplified by the K14-*Eda* mouse model with a 5-fold increase in the number of ductal tips (17). In the embryonic gland, branch angles were relatively homogenous centering around 90 degrees suggesting a means to maximally avoid the parental duct (in case of lateral branching) or the sister tip (after bifurcation) thereby leading to an efficient ‘spread’ of the epithelial network in a yet unobstructed stroma. The non-stereotypy of the ductal trees was most evident in the diverse branch lengths, which, however, did not follow a random distribution but gravitated toward the shorter end with a tail of very long ones thereby generating highly heterogeneous epithelial networks. Despite these polymorphic branch patterns, on average branching was remarkably constant as shown by the invariant ratio of the total length of the epithelial tree to branch number, and hence in this respect remarkably ‘stereotypic’.

Molecular regulation of branching morphogenesis has been investigated extensively (11, 12). Many mouse models display impaired ductal morphogenesis, yet in many cases it remains unclear whether growth (size of the ductal tree) and/or branching (branch point frequency per ductal length) is affected, as few studies report these parameters rigorously. Also, stroma-free organoid assays have provided valuable insights (25, 42, 43), though to what extent these findings reflect *in vivo* behavior of the mammary epithelium is uncertain. The autocrine, growth-inhibitory function of Tgf-β1 on postnatal mammary morphogenesis is well-described (23, 24, 25), but its role during embryogenesis has remained elusive. Our *ex vivo* organ culture experiment showed that Tgf-β1 treatment decreased branching frequency. This occurred at a low concentration of Tgf-β1 which did not impair overall growth suggesting a specific effect on branching probability. In contrast, suppression of Tgf-β activity *ex vivo* promoted growth, but did not alter branching frequency. These data are in line with previous analysis of pubertal *Tgfb1*^+/-^ mice expressing ∼10% of wild type levels of Tgf-β1, which show accelerated ductal growth without any apparent change in branch patterning (24).

Collectively, these data imply that endogenous Tgf-β1 signaling mainly functions to limit growth. Yet, there may be a narrow physiological range of Tgf-β1 levels and/or local variation in its activity that contribute to the choice of branch initiation, as suggested by *in vitro* studies (45).

Multiple in vivo studies link the Fgf pathway to mammary morphogenesis. In a genetic mosaic model, *Fgfr2* null epithelial cells are outcompeted by the wild type cells in TEBs due to a proliferation defect, and conditional attenuation of Fgfr1/2 activity severely compromises ductal growth (18, 19, 20) establishing an important function in proliferation, but revealing less about contribution to branching. Our data indicate that exogenous Fgf10 enhanced not only growth, but also branching. On the other hand, attenuated Fgf activity decreased growth without impairing branch frequency. One possible explanation for this counterintuitive finding is that some other signaling pathway may compensate for the initiation of branch formation (but not growth, as shown *in vivo*) when Fgf activity is suppressed. Alternatively, proliferation and branch point formation may require different Fgf activity levels, and/or even distinct downstream pathways.

Studies from other branching organs suggest that tube elongation may be driven via multiple mechanisms including convergent extension, collective migration, and oriented cell divisions (46, 47, 48, 49). The PCP pathway regulates all these processes and has been implicated in tubule elongation in the kidney where mutations in the PCP components including *Vangl2* result in dilated ducts (47, 49). Growth of the mammary ductal network is undoubtedly fueled by the proliferative tips (19), and we have recently identified oriented cell motility as an important cellular mechanism driving ductal growth (Myllymäki et al., 2023). Our current data suggest that other mechanisms may also contribute to ductal elongation: a mere change in the ductal aspect ratio can lead to elongation, implying involvement of tissue rearrangements. Cells of the main duct were oriented perpendicular to the elongating main duct suggesting that this might be an active process, but testing this hypothesis must await the development of a culture set-up capable of capturing this phenomenon *ex vivo*. A recent study assessing the mammary phenotype of mice homozygous for the *Looptail* (*Lp*) allele encoding a dominant negative Vangl2 protein that interferes also with the related Vangl1 protein (50), reported that 30% of *Lp* mammary epithelia transplanted into cleared fat pads generated outgrowths with wide ducts (51). This phenotype was not observed in single conditional *Vangl1* or *Vangl2* mutants, but was shared with mice heterozygous for *Prickle2*, another PCP component (46), indicating that PCP is involved in ductal elongation also in the mammary gland. Our data show that *Vangl2* null embryos had shorter roots (main ducts), but without concurring increase in diameter, as one might expect if convergent extension was compromised.

Instead, branching frequency was increased. This may indicate that the cell movements/rearrangements necessary for elongation are only partially impaired, e.g. due to redundancy with the related Vangl1 (51). Alternatively, some other Vangl2-dependent processes such as directional migration might be impaired. Further studies are warranted to clarify the role of Vangl2/PCP in mammary ductal morphogenesis.

Thus far, lateral branching has been reported to occur only in developing lungs and mammary glands. In the latter, side branches emerge during the estrous cycle and early pregnancy under the influence of hormonal cues, mainly progesterone (13). Here, by using time-lapse imaging we unequivocally establish that side branching is the predominant mode of branching in the embryonic mammary gland though tip bifurcations are also common. Our data indicate that the proximity to neighboring tips, rather than branch points, spatially constrain the emergence of new lateral branches. Whether lateral branching takes place during puberty has been a matter of controversy.

Pubertal glands are characterized by long extending “primary” ducts that repeatedly split into two new ones by bifurcation. Additionally, shorter “secondary” branches stem from the primary ducts. Earlier research has commonly interpreted (many of) the shorter branches to result from side branching events, a conclusion supported by whole mounts and histological sections revealing the emergence of nascent branches from an existing duct, often close to its invading tip (52, 53, 54, 55). A more recent study concluded that pubertal branching occurs near exclusively by tip bifurcations (10). Recent advances in intravital time-lapse imaging of postnatal mammary glands (56) should facilitate the efforts to decipher whether lateral branching occurs also during the pubertal period or whether puberty forms an exception in the journey of the developing mammary gland.

In conclusion, we report here the use of quantitative methodologies producing baseline data on embryonic mammary gland branching morphogenesis. The epithelial tree forms by a combination of side branching and tip bifurcation events (and to a small extent via trifurcations), similar to the developing lung (5). In contrast to the lung, however, rotations between branching events are minimal leading to the formation of a planar epithelial network from early on. In embryonic mammary glands, branch points form seemingly stochastically leading to highly variable network topologies, yet development progresses predictably owing to constant average branching frequency. How growth is “measured” even when branching is stochastic and how at the cellular level different signaling pathways modulate the branching frequency remain fascinating challenges for future research.

## Materials and methods

### Mice

The following mouse strains were used: ROSAmT/mG (Jackson Laboratory stock no. 007576) (57) (hereafter mT/mG mice) in a mixed C129SvJ/ICR background, and K14-Cre43 (hereafter K14-Cre) (58) and Fucci (RIKEN Tsukuba BioResource Center) (37) mice in an outbred NMRI background (Janvier Labs). *Vangl2* null allele in C57Bl/6 background was derived by serially crossing *Vangl2* ex2-3 floxed mice (Jackson Laboratory stock no. 025174) (59) with PGK-Cre mice universally expressing Cre (Jackson Laboratory stock no. 020811) (60) to permanently delete exons 2 and 3. All *Vangl2* KO embryos displayed severe neural tube defects confirming successful deletion (59). *Vangl2* null allele was genotyped with 5’-CATTTGTCTGTTGCTGGGTAA-3’ and 5’- TGGACTGTGTCTTGCCTACTG-3’ primers. The appearance of the vaginal plug was considered as embryonic day (E) 0.5. The embryonic stage was further verified by limb morphogenesis and other external criteria.

### Ethics statement

All mouse experiments were approved by the local ethics committee and the National Animal Experiment Board of Finland (licenses KEK16-021, KEK19-019 and ESAVI/2363/04.10.07/2017). Mice were euthanized with CO_2_ followed by cervical dislocation.

### Optical projection tomography

E16.5, E17.5, and E18.5 K14-Cre;mT/mG female embryos were decapitated and bisected and internal organs removed prior to fixation (4% PFA in PBS overnight at +4ᵒC). Individual mammary glands were dissected with a small amount of surrounding skin under the fluorescent microscope and immunohistochemically stained as previously described (61). Chicken IgY anti-GFP (GFP- 1010, 1:500, Aves Labs, Inc.) was used as the primary antibody, and Cy™3 AffiniPure Goat Anti- Chicken IgY (103-165-155, Jackson ImmunoResearch Europe Ltd.) as the secondary antibody.

Samples were mounted in SeaPlaque agarose (#50100, Lonza), dehydrated with methanol, cleared with BABB solution as described (61), and scanned with Bioptonics OPT Scanner 3001M with 6.7 µm camera pixel size and rotation step of 0.900. Scanned images were reconstructed with NRecon (Bruker Corp.). Mammary gland epithelium was roughly segmented by ImageJ (TrakEM package) and three-dimensional analysis was done with Tree Surveyor software (31). Maximum diameter was determined for each sample and skeletonization errors were manually corrected. In Tree Surveyor software all branching points were assigned as bifurcations. If two branches seemingly originated from the same branching point (segment lengths <7 µm), they were assigned as a trifurcation to avoid misconfiguration of branching generations. Terminal branches (tips) with a median diameter value=0 µm were considered artifacts and excluded from the analysis. The average diameter of the gland was calculated by formula 2 * √([total volume]/(π * [total length]). The average diameter of the main duct was calculated with the same formula with [volume of main duct] and [length of main duct]. For branch point angles, the local branch angle values were reported.

### Immunofluorescence and confocal microscopy

For whole-mount immunofluorescence staining, the ventral skin containing mammary glands was dissected from female E16.5, E17.5, or E18.5 mouse embryos. The samples were fixed in 4% PFA at 4 °C overnight and then washed with PBS for 3–4 hours. After permeabilization with 0.3% PBST (0.3% TritonX-100 in PBS) for 1–2 hours at room temperature, E16.5 and E17.5 samples were blocked with blocking buffer (5% normal donkey or goat serum, 0.5% BSA, and 0.5% TritonX-100 in PBS) for 1 h. E18.5 samples were blocked without permeabilization step with blocking buffer (5% goat serum, 1% BSA, and 0.3% TritonX-100 in PBS) for 1 h. All samples were incubated at 4°C with rat anti-mouse CD326 (EpCAM, BD Pharmingen, 552370, 1:1,000) and 10 μg/ml Hoechst 33342 (Molecular Probes/Invitrogen) in a blocking buffer for 2-3 days. After washing with 0.3% PBST for 3-4 hours, EpCAM was detected with an Alexa Fluor 488 or Alexa Fluor 647 -conjugated secondary antibody (1:500, Molecular Probes/Invitrogen) at 4°C for 2 days. For E18.5 samples, Phalloidin-Alexa 568 was included (1:300, Invitrogen) to visualize filamentous actin. After washing in 0.3% PBST for 2-3 hours, individual mammary glands with limited surrounding mesenchyme were dissected in PBS under a stereomicroscope and mounted within Vectashield (Vector Laboratories). The images were acquired in 1.10 µm (Fucci samples) or 1.40 µm (*Vangl2* null and wild type littermate samples) intervals by using a Zeiss LSM700 laser scanning confocal microscope with Plan-Apochromat 25x/0.8 immersion objective (Carl Zeiss Microscopy GmbH).

E18.5 *Vangl2* KO and their littermate control samples were cleared in a solution containing sucrose and glycerol overnight, mounted and images of main ducts were acquired with Leica Sp8 upright confocal microscope at 1.04 µm intervals with HC PL APO 20x/0.75 IMM CORR CS2 objective with glycerol.

### Organ culture and time-lapse imaging

Mammary buds were dissected from E13.5 embryos and cultured in Trowell-type culture setting at the liquid-air interface as previously described (29, 62). All embryos of the preferred genotype were used since both females and males develop similarly until E14.0. After dissecting the ventrolateral skin encompassing the mammary area, explants were treated for 25-35 min for Dispase II (Roche, 1.25U/ml) followed by 4-8 min with 0.75% pancreatin and 0.25% trypsin (Sigma-Aldrich) at room temperature. After a 30-40 mins incubation on ice in DMEM supplemented with 10% (vol/vol) FCS, 2 mM L-glutamine, and 20 U/ml penicillin-streptomycin (Gibco/Invitrogen), excess epithelium surrounding the mammary buds was microsurgically peeled off resulting in explants with mammary primordia attached to ample amount of mesenchyme. Mammary buds 2 and/or 3 were used for quantification in growth factor experiments. The culture medium consisted of a 1:1 mixture of DMEM with 2 mM l-glutamine and F12 (Ham’s Nutrient Mix) (both from Life Technologies) and was supplemented with 10% (vol/vol) FCS (HyClone/Thermo Scientific), 20 U/ml penicillin-streptomycin (Gibco/Invitrogen) and ascorbic acid (100 mg/L). Explants were cultured at 37°C with 5% CO_2_.

To test the effect of growth factors and signaling pathway inhibitors, mammary glands from wild type E13.5 NMRI embryos were cultured as described above. The following proteins and inhibitors were reconstituted according to the manufacturer’s recommendations and used at the indicated final concentrations: Fgf10 (R&D, 345-FG, 100ng/ml with 2 µg/ml heparin), SU5402 (CALBIOCHEM, 572630, 2.5 µM), Tgf-β1 (R&D, 7666-MB, 2.5 ng/ml), and SB431542 (TOCRIS Bioscience, 301836-41-9K, 10 µM). Cultured mammary glands begin branching on day 3, reflecting the in vivo onset of branching (29). To specifically assess the effect of the signaling molecules/inhibitors on branching rather than initial outgrowth, vehicle (untreated control; 2 µg/ml heparin in 0.1% BSA for Fgf10 and 4mM HCL in 0.1% BSA in PBS for Tgf-β1) or proteins/inhibitors were added on day 2 of culture; medium with the vehicle, protein, or inhibitor was changed every other day. From the second day of culture, explants were imaged once a day with a Zeiss Lumar microscope equipped with Apolumar S 1.2x objective.

Multi-position, automated time-lapse imaging of K14-Cre;mT/mG explants was used to assess the types of branching events (lateral branching, bi/trifurcations). Explants were dissected as described above, except that each explant had only mammary gland 3 to avoid any potential interference by the closely positioned mammary gland 2. Explants were placed on Transwell inserts (Costar, Corning Inc.) and cultured on 6-well plates allowing multiposition imaging. From day 3 to day 7 of culture, ductal trees were imaged with 3I Marianas widefield microscope equipped with 10x/0.30 EC Plan-Neofluar Ph1 WD=5.2 M27 at 37°C with 6% CO2. The medium was changed on day 2, at the onset of imaging (day 3), and once during imaging, on day 5. Images were acquired with a LED light source (CoolLED pE2 with 490 nm/550 nm) every 4 h.

### Image analysis

#### K14-Cre;mT/mG mammary gland explants

The types of branching events were assessed by careful examination of subsequent time point images. Because of 2-dimensional imaging, not all parts of the glands were in focus at all time points and occasionally the type of branching remained uncertain. Quantification of branching types was done only in glands in which at least 60% of branching events could be reliably determined. Time-lapse images were analyzed with ImageJ. Time series were registered with StackReg plugin (63) subtract background (rolling ball r=200) and Gaussian (sigma=2) filtered and binarized with the local threshold (Li method). Segmentation errors were manually corrected (i.e. gaps in mammary epithelium and separation of falsely connected branches), fill holes function and Shape Smoothing plugin were applied. Segmented image was skeletonized and skeletonization errors were manually corrected. Analyze Skeleton in 2D time series script (https://imagej.net/Analyze_Skeleton_2D_time_series) was used for measuring the skeletons. Epithelial area was measured from segmented images by ROI Manager in ImageJ and average ductal diameter was derived by formula [total area]/[total length] for each time point.

Ductal diameter prior to bifurcation was measured by fitting an ellipse perpendicular to the spline of the duct in the ductal end in the time point preceding the bifurcation event and measuring the major axis of the ellipse. Distance to the closest tip (bifurcations and lateral branching) and closest ipsilateral tip (lateral branching only) was measured using the “Straight line” tool and ROI Manager in ImageJ. Distance to neighboring branch points and length of the parental segment was measured from the skeletonized images in ImageJ. The minimum ductal diameter of the parental segment after the branching event was approximated by fitting a “Rotated Rectangle” to the narrowest point in the segmented images. The diameter was measured from each time point as three independent replicates which were inside a 2% difference. Measurements were not performed on branches forming next to each other at the same time point.

Ductal elongation measurement was done with Image J (Fiji) by using the plugin Analyze-skeleton- Analyze skeleton 2D/3D. The Excel file with branch length information was saved and x and y position of the fragment of interest from the tagged skeleton image was found and the branch length value corresponding to the x and y value was noted.

#### Cell cycle status quantification

To quantify cell cycle dynamics of epithelial cells in the developing mammary gland, E16.5 female embryos expressing Fucci reporters were stained with EpCAM as described above to discriminate the mammary epithelium and mesenchyme. The acquired confocal images were analyzed manually with Zen software (V2.3, Carl Zeiss Microscopy GmbH). Briefly, optical sections were selected every 10-11 µm in Z-axis, and mKO2 (cells in G1/G0) or mAG (cells in S/G2/M) expressing mammary epithelial cells (EpCAM+) were counted manually in selected optical sections. The root area was manually separated in Imaris at the position where the first branching occurs, and cells were counted accordingly.

#### Vangl2 knockout analysis

EpCAM whole-mount immunostained confocal images were processed with Imaris software for the 3D rendering of the mammary epithelium to visualize the ductal tree. The number of tips was counted manually based on the 3D view. Root analysis was done with the combined use of both Fiji ImageJ and Imaris software. The root area was manually separated as above. Masked stacks were made based on EpCAM staining, skeletonized with Fiji, and root length was quantified with the plugin script from (64). Root volume was quantified from 3D rendered images in Imaris. The average root diameter was calculated by formula 2 * √([total volume]/(π * [total length]).

For cell shape/volume analysis, the confocal image stacks were preprocessed with the ImageJ plugin Noise2Void (65) with the N2V train and predict module to remove the background. The training was performed with 100 epochs, 200 steps per epoch, a batch size per step of 64, and a patch shape of 64. The neighborhood radius was adjusted to either 4 or 5, based on the quality of the images. Cells were then segmented in 3D using a cell membrane marker, EpCAM, with Imaris software version 10 (Bitplane). Segmentation was manually examined and poor-quality cells were removed from later analysis. Cell volume was obtained using Imaris. To assess cell orientation, a Reference Frame was manually adjusted to align with the long axis of the main duct for each sample. The vector presents the cell elongation direction (ellipsoid axis length C) and the coordinates of the cell center were extracted for analysis in R with package “circular” (66) and “tidyverse” (67) . To measure the width of the main duct near the nipple and branch, we used the spot module of the Imaris software version 10 (Bitplane). Two spots were placed in 3D and the width of the duct was derived from the distance between the two spots and exported from Imaris.

#### Effect of growth factors/inhibitors on ex vivo cultured mammary glands

Quantification for the number of branches, and total ductal length of the mammary gland were done with Image J (Fiji).

### Statistical analysis

All data were analysed by Prism 7 and 9 (GraphPad Software), or by circular package for R (66). Statistical tests used are indicated in figure legends. p-values < 0.05 were considered significant. Throughout the figure legends: *p < 0.05, **p<0.01; ***p < 0.001, ****p < 0.0001. Rayleigh’s Z- test was cells of the main duct are randomly oriented with respect to the main duct by quantifying the angle between the cells’ longest aspect with respect to the elongating main duct (0° =parallel to the main duct; 90°= perpendicular to the main duct). Watson’s U2 test was used to analyze the difference in angle distribution between the wild type and Vangl2 KO samples.

## Acknowledgements

We thank Ms. Raija Savolainen and Ms. Riikka Santalahti for excellent technical assistance, and all Mikkola lab members for stimulating discussions. Dr. Ian Smyth is acknowledged for sharing the Tree Surveyor software, and Dr. Jukka Jernvall for advice. The mouse studies were carried out with the support of the HiLIFE Laboratory Animal Centre Core Facility, University of Helsinki. Confocal and widefield microscopy was performed at the Light Microscopy Unit of the HiLIFE- Institute of Biotechnology, University of Helsinki, and optical projection tomography at the BioImaging unit of the University of Oulu.

## Author Contributions

**Conceptualization:** RL, JS, MLM; **Formal analysis:** JS, RL, SMM, QL; **Investigation:** RL, JS, SMM, QL, ET, BK; **Methodology:** RL, JS, SMM, QL, MV, SK, RPH, SJV, MM; **Project administration:** MLM; **Resources:** MLM, SJV; **Supervision:** MLM; **Visualization:** JS, RL, SMM, ET, QL; **Writing – original draft:** RL, JS, SMM, MLM; **Writing – review & editing**: JS, RL, SMM, QL, SK, SJV, MLM

## References

1. Lu P, Werb Z. Patterning mechanisms of branched organs. Science. 2008;322(5907):1506-9.

2. Goodwin K, Nelson CM. Branching morphogenesis. Development. 2020;147(10).

3. Lang C, Conrad L, Iber D. Organ-Specific Branching Morphogenesis. Front Cell Dev Biol. 2021;9:671402.

4. Myllymaki SM, Mikkola ML. Inductive signals in branching morphogenesis - lessons from mammary and salivary glands. Curr Opin Cell Biol. 2019;61:72–8.

5. Metzger RJ, Klein OD, Martin GR, Krasnow MA. The branching programme of mouse lung development. Nature. 2008;453(7196):745-50.

6. Short KM, Smyth IM. The contribution of branching morphogenesis to kidney development and disease. Nat Rev Nephrol. 2016;12(12):754–67.

7. Short KM, Combes AN, Lefevre J, Ju AL, Georgas KM, Lamberton T, et al. Global quantification of tissue dynamics in the developing mouse kidney. Dev Cell. 2014;29(2):188–202.

8. Hauser BR, Hoffman MP. Regulatory Mechanisms Driving Salivary Gland Organogenesis. Curr Top Dev Biol. 2015;115:111–30.

9. Macias H, Hinck L. Mammary gland development. Wiley Interdiscip Rev Dev Biol. 2012;1(4):533–57.

10. Scheele CL, Hannezo E, Muraro MJ, Zomer A, Langedijk NS, van Oudenaarden A, et al. Identity and dynamics of mammary stem cells during branching morphogenesis. Nature. 2017;542(7641):313-7.

11. Spina E, Cowin P. Embryonic mammary gland development. Semin Cell Dev Biol. 2021;114:83–92.

12. Paine IS, Lewis MT. The Terminal End Bud: the Little Engine that Could. J Mammary Gland Biol Neoplasia. 2017;22(2):93–108.

13. Brisken C, Ataca D. Endocrine hormones and local signals during the development of the mouse mammary gland. Wiley Interdiscip Rev Dev Biol. 2015;4(3):181–95.

14. Wang Y, Chaffee TS, LaRue RS, Huggins DN, Witschen PM, Ibrahim AM, et al. Tissue- resident macrophages promote extracellular matrix homeostasis in the mammary gland stroma of nulliparous mice. Elife. 2020;9.

15. Luetteke NC, Qiu TH, Fenton SE, Troyer KL, Riedel RF, Chang A, et al. Targeted inactivation of the EGF and amphiregulin genes reveals distinct roles for EGF receptor ligands in mouse mammary gland development. Development. 1999;126(12):2739–50.

16. Ciarloni L, Mallepell S, Brisken C. Amphiregulin is an essential mediator of estrogen receptor alpha function in mammary gland development. Proc Natl Acad Sci U S A. 2007;104(13):5455–60.

17. Voutilainen M, Lindfors PH, Lefebvre S, Ahtiainen L, Fliniaux I, Rysti E, et al. Ectodysplasin regulates hormone-independent mammary ductal morphogenesis via NF-kappaB. Proc Natl Acad Sci U S A. 2012;109(15):5744–9.

18. Parsa S, Ramasamy SK, De Langhe S, Gupte VV, Haigh JJ, Medina D, et al. Terminal end bud maintenance in mammary gland is dependent upon FGFR2b signaling. Dev Biol. 2008;317(1):121–31.

19. Lu P, Ewald AJ, Martin GR, Werb Z. Genetic mosaic analysis reveals FGF receptor 2 function in terminal end buds during mammary gland branching morphogenesis. Dev Biol. 2008;321(1):77–87.

20. Pond AC, Bin X, Batts T, Roarty K, Hilsenbeck S, Rosen JM. Fibroblast growth factor receptor signaling is essential for normal mammary gland development and stem cell function. Stem Cells. 2013;31(1):178–89.

21. Zhang X, Martinez D, Koledova Z, Qiao G, Streuli CH, Lu P. FGF ligands of the postnatal mammary stroma regulate distinct aspects of epithelial morphogenesis. Development. 2014;141(17):3352–62.

22. Mailleux AA, Spencer-Dene B, Dillon C, Ndiaye D, Savona-Baron C, Itoh N, et al. Role of FGF10/FGFR2b signaling during mammary gland development in the mouse embryo. Development. 2002;129(1):53–60.

23. Silberstein GB, Daniel CW. Reversible inhibition of mammary gland growth by transforming growth factor-beta. Science. 1987;237(4812):291-3.

24. Ewan KB, Shyamala G, Ravani SA, Tang Y, Akhurst R, Wakefield L, et al. Latent transforming growth factor-beta activation in mammary gland: regulation by ovarian hormones affects ductal and alveolar proliferation. Am J Pathol. 2002;160(6):2081–93.

25. Nelson CM, Vanduijn MM, Inman JL, Fletcher DA, Bissell MJ. Tissue geometry determines sites of mammary branching morphogenesis in organotypic cultures. Science. 2006;314(5797):298-300.

26. Shamir ER, Ewald AJ. Three-dimensional organotypic culture: experimental models of mammalian biology and disease. Nat Rev Mol Cell Biol. 2014;15(10):647–64.

27. Hume RD, Pensa S, Brown EJ, Kreuzaler PA, Hitchcock J, Husmann A, et al. Tumour cell invasiveness and response to chemotherapeutics in adipocyte invested 3D engineered anisotropic collagen scaffolds. Sci Rep. 2018;8(1):12658.

28. Andl T, Reddy ST, Gaddapara T, Millar SE. WNT signals are required for the initiation of hair follicle development. Dev Cell. 2002;2(5):643–53.

29. Voutilainen M, Lindfors PH, Mikkola ML. Protocol: ex vivo culture of mouse embryonic mammary buds. J Mammary Gland Biol Neoplasia. 2013;18(2):239–45.

30. Veltmaat JM, Mailleux AA, Thiery JP, Bellusci S. Mouse embryonic mammogenesis as a model for the molecular regulation of pattern formation. Differentiation. 2003;71(1):1–17.

31. Short K, Hodson M, Smyth I. Spatial mapping and quantification of developmental branching morphogenesis. Development. 2013;140(2):471–8.

32. Speroni L, Voutilainen M, Mikkola ML, Klager SA, Schaeberle CM, Sonnenschein C, et al. New insights into fetal mammary gland morphogenesis: differential effects of natural and environmental estrogens. Sci Rep. 2017;7:40806.

33. Hannezo E, Scheele C, Moad M, Drogo N, Heer R, Sampogna RV, et al. A Unifying Theory of Branching Morphogenesis. Cell. 2017;171(1):242–55 e27.

34. Paine I, Chauviere A, Landua J, Sreekumar A, Cristini V, Rosen J, et al. A Geometrically- Constrained Mathematical Model of Mammary Gland Ductal Elongation Reveals Novel Cellular Dynamics within the Terminal End Bud. Plos Comput Biol. 2016;12(4).

35. Nerger BA, Jaslove JM, Elashal HE, Mao S, Kosmrlj A, Link AJ, et al. Local accumulation of extracellular matrix regulates global morphogenetic patterning in the developing mammary gland. Curr Biol. 2021;31(9):1903–17 e6.

36. Moses HL, Roberts AB, Derynck R. The Discovery and Early Days of TGF-beta: A Historical Perspective. Cold Spring Harb Perspect Biol. 2016;8(7).

37. Sakaue-Sawano A, Kurokawa H, Morimura T, Hanyu A, Hama H, Osawa H, et al. Visualizing spatiotemporal dynamics of multicellular cell-cycle progression. Cell. 2008;132(3):487–98.

38. Butler MT, Wallingford JB. Planar cell polarity in development and disease. Nat Rev Mol Cell Biol. 2017;18(6):375–88.

39. Belmonte JM, Swat MH, Glazier JA. Filopodial-Tension Model of Convergent-Extension of Tissues. Plos Comput Biol. 2016;12(6):e1004952.

40. Hadsell DL, Hadsell LA, Olea W, Rijnkels M, Creighton CJ, Smyth I, et al. In-silico QTL mapping of postpubertal mammary ductal development in the mouse uncovers potential human breast cancer risk loci. Mamm Genome. 2015;26(1-2):57–79.

41. Naylor MJ, Ormandy CJ. Mouse strain-specific patterns of mammary epithelial ductal side branching are elicited by stromal factors. Dev Dyn. 2002;225(1):100–5.

42. Goodwin K, Nelson CM. Uncovering cellular networks in branching morphogenesis using single-cell transcriptomics. Curr Top Dev Biol. 2021;143:239–80.

43. Huebner RJ, Neumann NM, Ewald AJ. Mammary epithelial tubes elongate through MAPK- dependent coordination of cell migration. Development. 2016;143(6):983–93.

44. Pierce DF, Jr., Johnson MD, Matsui Y, Robinson SD, Gold LI, Purchio AF, et al. Inhibition of mammary duct development but not alveolar outgrowth during pregnancy in transgenic mice expressing active TGF-beta 1. Genes Dev. 1993;7(12A):2308–17.

45. Pavlovich AL, Boghaert E, Nelson CM. Mammary branch initiation and extension are inhibited by separate pathways downstream of TGFbeta in culture. Exp Cell Res. 2011;317(13):1872–84.

46. Varner VD, Nelson CM. Cellular and physical mechanisms of branching morphogenesis. Development. 2014;141(14):2750–9.

47. Torban E, Sokol SY. Planar cell polarity pathway in kidney development, function and disease. Nat Rev Nephrol. 2021;17(6):369–85.

48. Tang Z, Hu Y, Wang Z, Jiang K, Zhan C, Marshall WF, et al. Mechanical Forces Program the Orientation of Cell Division during Airway Tube Morphogenesis. Dev Cell. 2018;44(3):313–25 e5.

49. Kunimoto K, Bayly RD, Vladar EK, Vonderfecht T, Gallagher AR, Axelrod JD. Disruption of Core Planar Cell Polarity Signaling Regulates Renal Tubule Morphogenesis but Is Not Cystogenic. Curr Biol. 2017;27(20):3120–31 e4.

50. Yin HF, Copley CO, Goodrich LV, Deans MR. Comparison of Phenotypes between Different vangl2 Mutants Demonstrates Dominant Effects of the Looptail Mutation during Hair Cell Development. Plos One. 2012;7(2).

51. Smith P, Godde N, Rubio S, Tekeste M, Vladar EK, Axelrod JD, et al. VANGL2 regulates luminal epithelial organization and cell turnover in the mammary gland. Sci Rep. 2019;9(1):7079.

52. Fata JE, Werb Z, Bissell MJ. Regulation of mammary gland branching morphogenesis by the extracellular matrix and its remodeling enzymes. Breast Cancer Res. 2004;6(1):1–11.

53. Silberstein GB. Postnatal mammary gland morphogenesis. Microsc Res Tech. 2001;52(2):155–62.

54. Silberstein GB, Daniel CW. Glycosaminoglycans in the basal lamina and extracellular matrix of the developing mouse mammary duct. Dev Biol. 1982;90(1):215–22.

55. Sternlicht MD, Kouros-Mehr H, Lu P, Werb Z. Hormonal and local control of mammary branching morphogenesis. Differentiation. 2006;74(7):365–81.

56. Lloyd-Lewis B. Multidimensional Imaging of Mammary Gland Development: A Window Into Breast Form and Function. Frontiers in Cell and Developmental Biology. 2020;8.

57. Muzumdar MD, Tasic B, Miyamichi K, Li L, Luo L. A global double-fluorescent Cre reporter mouse. Genesis. 2007;45(9):593–605.

58. Andl T, Ahn K, Kairo A, Chu EY, Wine-Lee L, Reddy ST, et al. Epithelial Bmpr1a regulates differentiation and proliferation in postnatal hair follicles and is essential for tooth development. Development. 2004;131(10):2257–68.

59. Copley CO, Duncan JS, Liu C, Cheng H, Deans MR. Postnatal refinement of auditory hair cell planar polarity deficits occurs in the absence of Vangl2. J Neurosci. 2013;33(35):14001–16.

60. Lallemand Y, Luria V, Haffner-Krausz R, Lonai P. Maternally expressed PGK-Cre transgene as a tool for early and uniform activation of the Cre site-specific recombinase. Transgenic Res. 1998;7(2):105–12.

61. Alanentalo T, Asayesh A, Morrison H, Loren CE, Holmberg D, Sharpe J, et al. Tomographic molecular imaging and 3D quantification within adult mouse organs. Nat Methods. 2007;4(1):31–3.

62. Lan Q, Satta J, Myllymaki SM, Trela E, Lindstrom R, Kaczynska B, et al. Protocol for Studying Embryonic Mammary Gland Branching Morphogenesis Ex Vivo. Methods Mol Biol. 2022;2471:1–18.

63. Thevenaz P, Ruttimann UE, Unser M. A pyramid approach to subpixel registration based on intensity. IEEE Trans Image Process. 1998;7(1):27–41.

64. Arganda-Carreras I, Fernandez-Gonzalez R, Munoz-Barrutia A, Ortiz-De-Solorzano C. 3D reconstruction of histological sections: Application to mammary gland tissue. Microsc Res Tech. 2010;73(11):1019–29.

65. Broaddus C, Krull A, Weigert M, Schmidt U, Myers G. Removing Structured Noise with Self- Supervised Blind-Spot Networks. I S Biomed Imaging. 2020:159–63.

66. Agostinelli C, Lund U. R package ‘circular’: circular statistics (version 0.4-93). URL https://r-forger-project.org/projects/circular. 2017.

67. Wickham H, Averick M, Bryan J, Chang W, McGowan LDA, François R, et al. Welcome to the Tidyverse. Journal of open source software. 2019;4(43):1686.

